# Identifying *Streptococcus pneumoniae* genes associated with invasive disease using pangenome-based whole genome sequence typing

**DOI:** 10.1101/314666

**Authors:** Uri Obolski, Andrea Gori, José Lourenço, Craig Thompson, Robin Thompson, Neil French, Robert Heyderman, Sunetra Gupta

**Affiliations:** University of Oxford, Department of Zoology, Oxford, UK; University College London, Division of infection and immunity, London, UK; Liverpool School of Tropical Medicine, Liverpool, UK

## Abstract

*Streptococcus pneumoniae* is a normal commensal of the upper respiratory tract but can also invade the bloodstream or CSF (cerebrospinal fluid), causing invasive pneumococcal disease (IPD). In this study, we attempt to identify genes associated with IPD by applying a random forest machine-learning algorithm to whole genome sequence (WGS) data. We find 43 genes consistently associated with IPD across three geographically distinct WGS data sets of pneumococcal carriage isolates. Of these genes, 23 genes have previously shown to be directly relevant to IPD, while the other 18 are uncharacterized.

## Introduction

Invasive pneumococcal disease (IPD) is defined as an infection in which the bacterial pathogen *Streptococcus pneumoniae* (pneumococcus) enters a usually sterile site, such as the blood or cerebrospinal fluid [1]. Although pneumococci are usually carried asymptomatically within the human nasopharynx, IPD is life-threatening and constitutes a major cause of mortality, disproportionally affecting children, elderly and immunocompromised individuals [2, 3]. Genetic changes facilitating the survival of pneumococci during invasion have been previously identified and described through experimental and bioinformatic methods [4–10]. The work of Hava and Cammilli, for instance, describes a set of 387 transposon insertion mutants exhibiting attenuated virulence in mice in the model pneumococcal strain TIGR4, and defines the genes disrupted in each mutant as putative virulence factors [4]. Several other studies have been successful in identifying differential expression patterns of key virulence genes of *S. pneumoniae* in vitro and in vivo [5, 6]. These studies used RT-PCR on previously described virulence factors and high-throughput microarray expression profiling to identify gene expression signatures during invasion of model organisms or growth on epithelial cell lines. DNA microarrays have also been employed in order to identify a common core genome differentiating between strains isolated from invasive disease or carriage in three pneumococcal serotypes often found in IPD (6A, 6B and 14) [8]. Although these methods did highlight features involved in the ability of pneumococci to invade a host, they were limited either by small sample size, representing only on a fraction of the pneumococcal serotypes, or by relying on a single reference genome to identify patterns of differential gene expression and gene presence in IPD isolates. Recent studies which used whole-genome sequence data failed to identify adaptive differences, in terms of presence and absence of genes or genetic mutations, between strains invading the blood and strains that were able to cross the blood-brain barrier [11–13]. These studies highlight the need of future research to comprehensively identify whether adaptation of IPD isolates occurs through genetic variation between carriage and invasion, suggesting that subtle changes may influence the virulence of the bacterial isolates.

Here, we looked for genetic differences between pneumococcal carriage and IPD isolates using a whole genome sequence typing approach. In this approach, every gene in a data set is assigned an allelic coding based on a designated reference genome, and new alleles are defined as gene variants containing any change from previously defined alleles (see Methods). This is an extension of the widely used multilocus sequence typing (MLST) method [14, 15]. However, in contrast to MLST, which is based on a small number of genes usually present in all isolates of a bacterial species, sequence typing of an entire bacterial genome may contain substantial variations in gene presence.

Hence, using a particular reference genome for such an analysis might preclude identification of genes not found in a certain serotype. To overcome this problem, we constructed a pangenome of our IPD samples and used it as a synthetic reference genome. We then employed the random forest algorithm (RFA) to identify which genes are the best predictors of invasive/carriage phenotypes; this machine learning algorithm is commonly applied to genomic data [16] and has been previously used by our group identify that genes under immune selection were associated with the lineage structure of pneumococci [17]. To reduce the confounding effect of bacterial lineages structure, we performed this analysis on three data sets, each time complementing the IPD isolates with carriage isolates from a different country (UK, USA and Iceland). We found 43 genes in all three comparisons to rank among the top most predictive genes for IPD, many of which were supported in the literature as associated with IPD. Our method was also compared to an approach classifying carriage/invasiveness based solely on presence-absence of genes as well as to a WGS maximum likelihood tree-based approach, clustering isolates to clades. Finally, we analyzed the identified genes’ length and location on the pneumococcal genome relative to capsule-determining loci.

## Results

We obtained 378 pneumococcal isolates causing bacteremia, from different countries, as presented in Table S1. The number of invasive isolates was limited by public availability of WGS samples marked as isolated from blood. A pangenome of 9032 genes was generated from this data set, from which all genes in the soft-accessory genome (defined as genes appearing in at least 15% of samples) were used as a reference genome for the sequence typing process. The sequence typing process was applied three times: on the invasive disease isolates (n=378) joined with a data set of carriage isolates from the UK (n=520), USA (n=622) and Iceland (n=622). The three datasets were not combined to a single data set for two reasons: first, comparing the results from three different countries constitutes a more conservative approach, increasing the probability of finding genes truly associated with invasive disease rather than associated with lineages more prevalent in certain datasets; Second, the computational complexity of sequence typing increases non-linearly with the number of genomes and prevents large datasets from being typed simultaneously. Since typing (i.e. the ‘naming of alleles’) is arbitrary, separately typed datasets cannot be combined. Therefore, we used all carriage isolates available from the UK, and as many isolates from Iceland and the US as possible while maintaining a total of no more than 1000 sequences (the limit in BIGSdb, the web server used for the typing service – see Methods).

RFA was applied to each of these data sets with invasive/non-invasive disease as the predicted variable. The out-of-bag (OOB) classification success [18] was similar for the three datasets with 94.6% (95% CI 94.6-94.7), 93.2% (95% CI 93.1-93.2), and 94.4% (95% CI 94.4-94.5) success for carriage, and 76.7% (95% CI 76.7-76.8), 86.7% (95% CI 86.7-86.8), and 79.7% (95% CI 79.6-79.8) success for bacteremia in Iceland, the UK, and USA, respectively. For each data set, the top 100 genes with the highest importance score were chosen, using a heuristic method aiming to maximize the fraction of joint genes (see Methods), and then recorded and compared. Out of these, 43 were joint to all three data sets. All the identified genes are presented in Table 1. The probability of this many, or more, genes joint to all three data sets in a random selection of 100 genes (i.e. the p-value for the null hypothesis of the RFA choosing genes randomly) was verified via simulations to be <10-^6^. Furthermore, RFA was run again using only the 43 genes joint to the three data sets. The OOB classification success was 93.2% (95% CI 93.1-93.2), 96% (95% CI 95.9-96), and 95.9% (95% CI 95.9 -96) for carriage, and 73.4% (95% CI 73.3-73.4), 90.1% (95% CI 90- 90.2), and 80.6% (95% CI 80.4-80.7) success for bacteremia, in Iceland, the UK, and USA, respectively. This indicates that the 43 joint genes are providing sufficient information to classify invasive versus non-invasive pneumococcal strains.

**Table 1:** Genes associated with IPD.

| Gene | Length (bp) | Best matches, identity (%), e-value, Accession number | Description |
| --- | --- | --- | --- |
| *hpp1* | 452 | 1. phtB, 9E-132, 447/465 (96%), NCBI, AF318954.1<br>2. phpA, 0, 451/480 (94%), NCBI, AF340221.1 | 1. PHT proteins (aka BHV) are thought to be involved in the invasion process of pneumococci [23, 24].<br>2. The PhpA protein elicits protective immune response against bacteremia and nasopharyngeal carriage in mice [23]. |
| *hpp2* | 330 | hypothetical protein (CPS), 4E-168, 328/330 (99%), NCBI, JQ653094.1 | This is a putative capsular polysaccharide biosynthesis protein, Capsular differences are known to be associated with invasive disease [7]. |
| *hpp3* | 249 | hypothetical protein |  |
| *hpp4* | 504 | phtD, 0, 504/504 (100%), NCBI, KP127799.1 | The found phtD hit was a part of a sequence shown to be highly conserved in invasive isolates [25]. |
| *hpp5* | 954 | Hypothetical protein (CPS), 0, 954/954 (100%) , NCBI, HE651314.1 | This is a putative capsular polysaccharide biosynthesis protein. Capsular differences are known to be associated with invasive disease [7]. |
| *hpp6* | 996 | Hypothetical protein |  |
| *hpp7* | 231 | pspC, 1E-67, 167/179 (93%), NCBI, AF154043.2 | pspC was shown to be involved in immune response to bacteremia in mice [26]. |
| *hpp8* | 510 | Hypothetical protein |  |
| *hpp9* | 324 | Hypothetical protein (CPS), 2E-161, 320/324 (99%), NCBI, ADM91299.1 | This is a putative capsular polysaccharide biosynthesis protein. Capsular differences are known to be associated with invasive disease [7]. |
| *hpp10* | 504 | pspC,0, 502/504 (99%), NCBI, AF154022.1 | pspC was shown to be involved in immune response to bacteremia in mice [26]. |
| *hpp11* | 306 | Hypothetical protein |  |
| *hpp12* | 399 | Hypothetical protein |  |
| *hpp13* | 327 | Hypothetical protein (CPS), 0, 511/528 (97%), NCBI, AF316639.1 | This is a putative capsular polysaccharide biosynthesis protein. |
|  |  |  | Capsular differences are known to be associated with invasive disease [7]. |
| ydcP_1 | 471 | putative protease YdcP, 0, 470/471(99%), NCBI, AFS43444.1 | YdcP is part of the U32 protease family. It is a collagenase, facilitating breaking of extracellular structures tissues, and is a known virulence factor in other bacterial species [27]. |
| *hpp14* | 519 | Hypothetical protein (CPS), 0 509/519 (98%), NCBI, AF154022.1 | This is a putative capsular polysaccharide biosynthesis protein. Capsular differences are known to be associated with invasive disease [7]. |
| *hpp15*<br>(*hmo*) | 147 | L-lactate dehydrogenase (FMN-dependent)-like /alpha-hydroxy acid dehydrogenase, 4E-70, 147/147(100%) | Lactate dehydrogenase was found to be essential enzyme for pneumococcal survival in blood [28]. |
| *hpp16* | 480 | Hypothetical protein |  |
| *lytB* | 1977 | Putative endo-beta-N-acetylglucosaminidase, 0, 1968/1977 (99%), NCBI, AJ870414.1 | *lytB* codes for a endo-beta-N-acetylglucosaminidase, which is responsible for cell-wall hydrolysis and is thought to be a virulence factor [29, 30]. |
| *hpp17* | 528 | Hypothetical protein (CPS), 511/528 (97%), NCBI, JF301964.1 | This is a putative capsular polysaccharide biosynthesis protein. Capsular differences are known to be associated with invasive disease [7]. |
| *hpp18* | 516 | pspC, 508/516 (98%), NCBI, AF154043.2 | pspC was shown to be involved in immune response to bacteremia in mice [26]. |
| *hpp19* | 489 | Hypothetical protein |  |
| *hpp20* | 387 | Hypothetical protein (partial transposase), 0, 387/387(100%), NCBI, ADM91518.1 | Part of the mobile genetic elements of the bacterium. |
| *hpp21* | 258 | Hypothetical protein |  |
| *hpp22* | 288 | Hypothetical protein |  |
| *hpp23* | 210 | Hypothetical protein |  |
| *hpp24* | 387 | Hypothetical protein (partial transposase), 0.0, 386/387(99%), NCBI, | Part of the mobile genetic elements of the bacterium. |
|  |  | CP002176 (positions 1374937-1375323) |  |
| *hpp25* | 510 | Hypothetical protein |  |
| *hpp26* | 168 | Hypothetical protein |  |
| *hpp27* | 489 | pspC, 0, 463/490 (94%), NCBI, AF154022.1 | pspC was shown to be involved in immune response to bacteremia in mice [26]. |
| *lox* | 1137 | Lactate oxidase (*lox*) gene, 0, 1001/1137(88%), NCBI, DQ984140.3 | The *lox* gene is involved in bacterial niche competition and virulence in streptococci and other bacterial species [31, 32]. |
| *hpp28* | 840 | Sortase (*srtA*), 0, 614/740(83%), NCBI, KX147105.1 | In *Streptococcus mutans*, disruption of the sortase (*srtA*) gene led to decrease in adherence and invasion to endothelial cells [33]. |
| *hpp29* | 189 | Hypothetical protein |  |
| *hpp30* | 537 | Hypothetical protein |  |
| *hpp31* | 504 | Hypothetical protein |  |
| *hpp32* | 309 | Hypothetical protein |  |
| *hpp33* | 309 | Hypothetical protein |  |
| *cpsA* | 1446 | *cpsA* (aka *wzg*), 0, 1446/1446 (100%), NCBI, KC522490.1 | *wzg* (aka *cpsA*) is part of the capsular polysaccharide synthesis gene locus. High expression of *cpsA* is associated with bacteremia in humans [34]. |
| *bgaA* | 6702 | *bgaA* (Beta-galactosidase BoGH2A), 6466/6704 (96%), NCBI, AF282987.1 | *bgaA* is hypothesized to be a pneumococcal virulence factor [35] and was shown to promote resistance to immune cells in human serum [36]. |
| *cpsA* | 1446 | *cpsA* (aka *wzg*), 0, 1446/1446 (100%), NCBI, KC522492.1 | *wzg* (aka *cpsA*) is part of the capsular polysaccharide synthesis gene locus. High expression of *cpsA* is associated with bacteremia in humans [34]. |
| *hpp34* | 207 | Hypothetical protein |  |
| *hpp35* | 573 | pspC, 0, 566/573 (99%), NCBI, AF154043.2 | pspC was shown to be involved in immune response to bacteremia in mice [26]. |
| *hpp36* | 684 | cpsD, 0, 682/684 (99%), NCBI, AFC94091.1 | cpsD mutations were shown to inhibit the possibility of causing bacteremia in mice [37]. |
| *hpp37* | 840 | Hypothetical protein |  |

We compared our method to two established analysis methods: first, we repeated the analysis based on a genome-wide presence and absence of genes, rather than their alleles, using Scoary [19]. No genes were identified as jointly highly predictive in all three data sets using Scoary, even when the top-300 ranking genes found by this method were considered. We then applied a sequence-based maximum likelihood phylogeny of the core genes of each dataset [20]. This method also could not capture the genetic differences between the invasive and carriage isolates, as these isolates remained scattered across different clades (see Methods and supplementary material Figures S3-S5).

Interestingly, 23 of the genes we identified had BLAST matches with genes previously found to be associated with invasive disease or associated with immune response to it. 18 genes were found to encode for hypothetical proteins with unknown functions and 2 genes were found to encode for transposases, which catalyse the rearrangement of mobile genetic elements in the bacterial chromosome [21]. We explored the characteristics of the identified IPD-associated genes by determining the locations of the identified genes on the pneumococcal genome. Figure. 1A shows the locations of the identified genes across the genome of a 19A serotype sample and reveals that they are spread across the pneumococcal genome. Several identified genes were found near this capsule polysaccharide synthesis locus (orange rectangle on Figure. 1A), but many of the other genes identified were spread across the genome, suggesting that our findings do not simply rely on differences in serotype compositions between the datasets used for our analysis (for serotype distribution in our data, see Figure S1). We note that since pneumococcal serotypes have substantial genomic variation, driven by recombination, horizontal gene transfer and events of gene loss or addition, the locations of genes within their respective genomes are not constant [22]. Regardless, qualitatively similar location distributions were obtained when plotting these genes on other serotype samples (SI Figures S5, S6). In addition to this, we examined the length of the IPD associated genes (Figure. 1B), since many of them were found to have BLAST matches to short subsets of known genes (see Table 1). The IPD associated genes were statistically significantly shorter than those in the soft-accessory genome (Wilcoxon rank-sum test, p-value = 0.00027) but not significantly shorter than those of the entire pangenome (Wilcoxon rank-sum test, p-value =0.079). Furthermore, the length of genes in the entire pangenome was significantly shorter than those in the soft-accessory genes (Wilcoxon rank-sum test, p-value = 10^-16^). I.e., the genes we identified are comparable in length to those in the entire pangenome, but are shorter than the soft-accessory genes we used as a reference. Thus it seems that shorter genes are associated with having a variable presence across pneumococci, and a higher probability of being associated with IPD.

**Figure 1:**
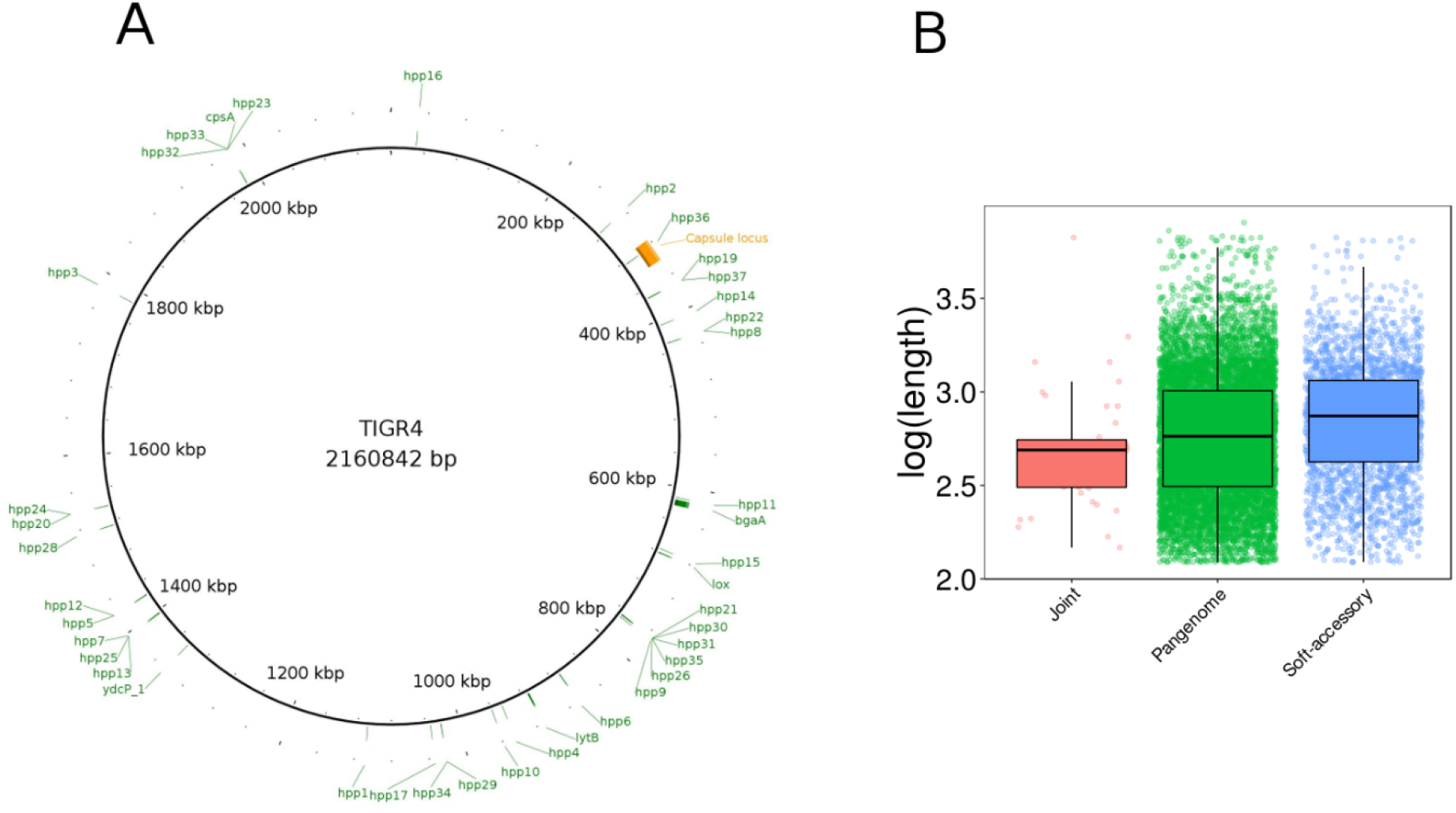
Location and length of genes associated with IPD. **(A)** Location of identified IPD-associated genes (see Table 1) on a 19A streptococcal genome (accession NC_010380.1). Orange rectangle marks the capsular synthesis locus (CPS). Similar plots using other serotype samples can be found in Figures S5-S6. **(B)** Boxplots and distributions of log10-transformed gene lengths from the IPD-associated genes, the entire pangenome and the soft-accessory genome used in our analysis (see methods).

## Discussion

In this study we identified pneumococcal genes associated with IPD using a novel method, comprising several techniques. First, we encoded WGS data by extending a multi-locus sequence typing scheme to the entire genome. This approach enables information to be extracted from gene variants, or alleles, as well as from the presence/absence of genes.

As a reference genome for the typing scheme, we constructed a genome which included any genes existing in more than 15% of invasive samples, namely the soft-accessory genome. This is especially important when typing pneumococcal samples, which have highly variable genomes and can yield a core genome shorter than 50% of an average pneumococcal genome [38, 39]. Using a reference genome constructed in such a way proved beneficial, as all but one of the genes eventually identified as associated with IPD were present in fewer than 95% of isolates, categorizing them in the soft-accessory genome rather than in the core genome (SI Table S2).

We then used an RFA to find the best predictors of invasive disease and carriage. RFA creates multiple decision trees, selecting for each tree a subset of genes and a bootstrap sample of the isolates. It than scores the genes by their marginal contribution to improving classification of invasive disease and carriage on samples not selected in the bootstrapping process. Our method was implemented on three datasets of pneumococcal carriage samples isolated from different countries, and the top-ranking genes were reduced to only those that were jointly top-ranking in all three datasets. Selecting the jointly top-ranking genes imposes a stringent cutoff for the identified genes, and reduces potential bias introduced due to local ancestry or population structure. It resulted in a total of 43 jointly high scoring genes out of 100 top-ranking genes associated with IPD – implying a relatively high replicability of results across datasets. For comparison, we applied a presence/absence method and a sequence-based phylogenetic approach, which yielded no significant results joint to all three data sets.

Reassuringly, many of the genes we identified are parts of known virulence factors, or are associated with invasive pneumococcal disease and especially with bacteremia (see Table 1). The gene *lytB*, for instance encodes the LytB protein which is involved in the attachment of *S. pneumoniae* to human nasopharyngeal cells in vitro. Loss of *lytB* has been shown to heavily impair pneumococcal virulence in a mouse sepsis model [29, 30]. It has also been shown that LytB is essential for a successful biofilm production and avoidance of phagocytosis [40]. Another gene we identified here was the lactate oxidase, *lox*. In other streptococcal species, particularly *S. mutans, S. pyogenes* and *S. oligofermentas*, H_2_O_2_- producing lactate oxidase activity was shown to be used in absence of glucose and for niche competition [31, 32]. *S. pneumoniae* is also known to use lactate as an energy source in absence of glucose, converting the lactate molecule to pyruvate with consequent production of H_2_O_2_ [41]. Although *S. pneumoniae* lacks the catalase enzyme, which plays a central role in bacterial resistance to hydrogen peroxide, it is able to detoxify the large quantities of H_2_O_2_ produced during colonization through mechanisms that have yet to be fully elucidated [42]. However, H_2_O_2_ production confers a colonization advantage in the upper respiratory tract due to its bactericidal effect on other colonizing bacteria [43]. Moreover, the H_2_O_2_ secreted by *S. pneumoniae* was shown to facilitate DNA damage, and cell apoptosis, contributing to bacterial virulence [44].

Four homologs of *pspC* (which is known to exhibit high copy-number and allelic variation [45, 46]) appear among the genes we identified, namely *hpp7, hpp10, hpp18 and hpp35*. PspC is a bacterial surface protein (adhesin) essential for colonization of nasal tissue [47]. The protein binds to endothelial blood-brain barrier receptors, potentially facilitating CNS invasion [48]. Moreover, PspC binds the human factor H, which is identified as a key mechanism to avoid complement deposition [49]. Recently, two subgroups of PspC have been identified (PspC I and PspC II), with the subgroup one being more represented in IPD isolates and more effective in complement evasion than subgroup two [49]. The repeated identification of several copies of PspC by our method endorses its proposed use in a non-capsular vaccine conferring protection against carriage and invasive disease in a mouse model [26, 47]. Even though the high variability of PspC was considered an obstacle for such a vaccine [50], our results can mitigate this shortfall by suggesting which variants of the protein might be more relevant for IPD. This can allow such a vaccine to have sufficient efficiency by targeting the most relevant variants of the protein, despite having a low coverage of all existing variants.

Two of the genes we found to associate with IPD encoding for transposases. It is known that *S. pneumoniae* is characterized by a high level of genomic plasticity, which allows to the bacterium to react quickly to changes in environmental conditions [51]. As mobile genetic elements are responsible for the dissemination of phenotypic characteristics in the bacterium, such as antimicrobial resistance [52], and are overexpressed in conditions related to virulence, such as during biofilm production [53], it is possible to speculate that these mobile genetic elements could be associated with the dissemination of virulence factors amongst the *S. pneumoniae* species.

Most of the other identified genes were hypothetical, with no known function. Based on our method’s classification success, the fact that the highly ranked genes were identified in the analyses of three independent carriage datasets, and the high presence of known virulence factors among the genes, we believe that the hypothetical genes identified are highly likely to be involved in pneumococcal invasive disease. Of particular interest are identified genes which are farther from the capsular locus (see Figure 1.A), which could potentially be serotype-independent IPD-associated genes and therefore relevant across pneumococcal strains. The length of the identified genes was also unusually short relative to the synthetic reference genome we used (Figure 1.B) suggesting that further attention should be given to shorter sequences and gene fragments as potential factors contributing to IPD.

The main limitation of our method is that all alleles are marked as different ‘states’ of a gene and their degree of similarity/difference is not taken into account. Thus, we can identify which genes are associated with differing phenotypes, but subtler methods will be necessary to discern exactly which alleles are responsible for which phenotypic changes. A feasible future extension of our method could be adding variables encoding more information about the alleles, such as structural properties of their resulting proteins [54]. However, such an extension will necessitate an efficient way of combining the genetic and protein information as the interactions between genes and their translated protein characteristics will likely have a substantial effect on the results.

Moreover, by restricting the genes we identify to only those that are highly ranked in multiple datasets our method trades sensitivity for specificity. For example, *prtA* is known to be associated with IPD [29], but was ranked as 96^th^, 121^st^, and 240^th^ in the US, UK and Iceland datasets, respectively. Hence, changing the cutoff threshold to 250 joint genes would have identified this gene as well. Other genes may be excluded from our results because they are not uniformly predictive of IPD across populations, due to the different genetic backgrounds of pneumococci or due to sampling noise in the available data. This is the case with *lytA*, which is known to be associated with IPD [7] and was ranked 129^th^ and 99^th^ in the US and Iceland datasets, but only 739^th^ in the UK dataset. Finally, genes necessary for an invasive phenotype might also not be found by our method if they are present and relatively conserved in all isolates, as was indeed the case with *ply* [25].

We believe the method presented here can be applied to a variety of pathogens to identify genes responsible for virulent phenotypes. We foresee our approach being particularly useful when the examined pathogens share only a small core genome, such as *E.coli* [55] and *C. jejuni* [56]. The goal of our method is to discern with high confidence genes associated with IPD, or any other phenotype, so their function could eventually be experimentally examined. Accordingly, we hope the hypothetical genes identified in this study will be further analyzed and prove to be useful in our understanding of invasive pneumococcal disease.

## Methods

### Pangenome construction and sequence typing

A total of 378 genome sequences of *S. pneumoniae* strains isolated from invasive disease were downloaded from BIGSdb [57] with geographical origin of isolates and accession numbers available in SI Table S1.

These genomes were used to build an invasive population pangenome using Roary V.3.6.1 [58]. Briefly, each draft genome downloaded from BIGSdb was re-annotated with PROKKA V1.12 [59] and the annotation output was fed to Roary for the pangenome construction. Roary parameters were set to minimum blastp identity 90% and MCL inflation value of 1.5. For the purpose of this analysis, we included in the pangenome the genes present in the soft-accessory genome, i.e. present in >15% of isolates, for a total of 2649 genes. This pangenome was used as the reference genome for sequence typing of three new datasets, containing the invasive sequences together with each one of the three carriage data sets, namely Iceland, the UK or USA. BIGSdb parameter values were set to the webserver defaults. Genome sequences were quality controlled before the pan-genome construction by making sure that the total length of each assembly was between 2.0 and 2.3 Mb (the common genome length of completed *S. pneumoniae* genomes as retrieved from https://www.ncbi.nlm.nih.gov/genome/). Moreover, the absence of low-level contamination was ascertained using Kraken v 0.10.5 [60]. Briefly, if more than 5% of the total genome assembly sequence was identified as belonging to a different bacterial species, that assembly was removed from further analysis. As shown in supplementary material figures S8 and S9, satisfactory pangenome saturation was reached in terms of core genome and number of new genes added per new genome [61].

### Scoary, FastTree and genome-location

A pangenome wide association study (Pan-GWAS) was performed using the Scoary V.1.6.16 pipeline [19]. Three new pan-genomes were built using 3 different datasets, each including the 378 genome sequences from the invasive disease strains and genomes from the carriage strains isolated from either Iceland, the UK or USA. Each of the three pan-genomes (invasive+carriage strains) was then input to Scoary using the invasive/carriage origin of the strain as classifier for the pan-GWAS pipeline. The genes representing the core genomes of the three invasive+carriage datasets (present in more than 99% of the analysed isolates) were concatenated and aligned with MAFFT V.7.221 [62]. The alignment of the core genome was used to reconstruct the maximum likelihood phylogeny of each group of isolates using FastTree V.2.1 [20] under a generalized time-reversible model. Phylogenetic trees for each datasets were then edited and annotated using Evolview V.2 [63]. Genome location plots were produced using BRIG V.0.95 [64] with the genome sequence of strains 19A (NC_010380.1), D39 (NC_008533.1) or R6 (NC_003098.1) as references (Figure. 1, Figure S5 and Figure S6, respectively).

### Random forest analysis

Random forest was implemented in R using the *randomForest* package V.4.6-12 [65]. Allele types were turned into numeric variables in the RFA due to computational limitations. To break any biases such enumerations might introduce, we permuted each allele typing and reran the RFA for 200 times on each dataset [17]. The measure used to rank genes was permutation importance (aka Breiman-Cutler importance). Under this method, variable values are permuted for the OOB data of each tree and the resulting classification error is subtracted from the OOB data error without the variable permutation [18]. The average of this difference across all trees is the permutation importance. These importance measures were ranked for all variables and the rankings were averaged across the 200 permutations of RFA applications on each dataset [17]. The fraction of genes joint to the three datasets was compared as a function of the number of top-ranked genes selected. To reduce noise due to small samples of top-ranked genes, both the fraction of genes and the low bound of a 95% binomial confidence interval (with n=number of top-ranked genes and p=fraction of joint genes) were used. In both measures, the maximum fraction corresponded to using 100 top-ranked genes (Figure S7). Although similar peaks occur when more top-ranked genes were used, we chose 100 as a conservative threshold (i.e. to reduce the number of false positive genes identified).

### Functional annotation of genes associated with IPD

All gene sequences in Table 1 were first functionally annotated using the NCBI conserved domain search engine (https://www.ncbi.nlm.nih.gov/Structure/cdd/wrpsb.cgi). Each DNA and translated amino acid sequence was checked for similarity against known genes and protein using nucleotide and protein blast (megablast and blastp algorithms respectively, https://blast.ncbi.nlm.nih.gov). The combined results of the conserved-domains search and blast are described in Table 1.

### Data availability

Accession numbers for pneumococcal sequences used are listed in SI Table S1; the pangenome built from invasive isolates can be found in SI Table S3.

## Acknowledgements

This research was supported by an EMBO postdoctoral fellowship (UO), the European Research Council under the European Union’s Seventh Framework Programme (FP7/2007- 2013) /ERC grant agreement no. 268904 – DIVERSITY (JL and SG), the Medical Research Council MR/N023129/1 (AG, NF, RH and SG), and a Junior Research Fellowship from Christ Church, Oxford (RT).

## Supplementary information

**Figure S1:**
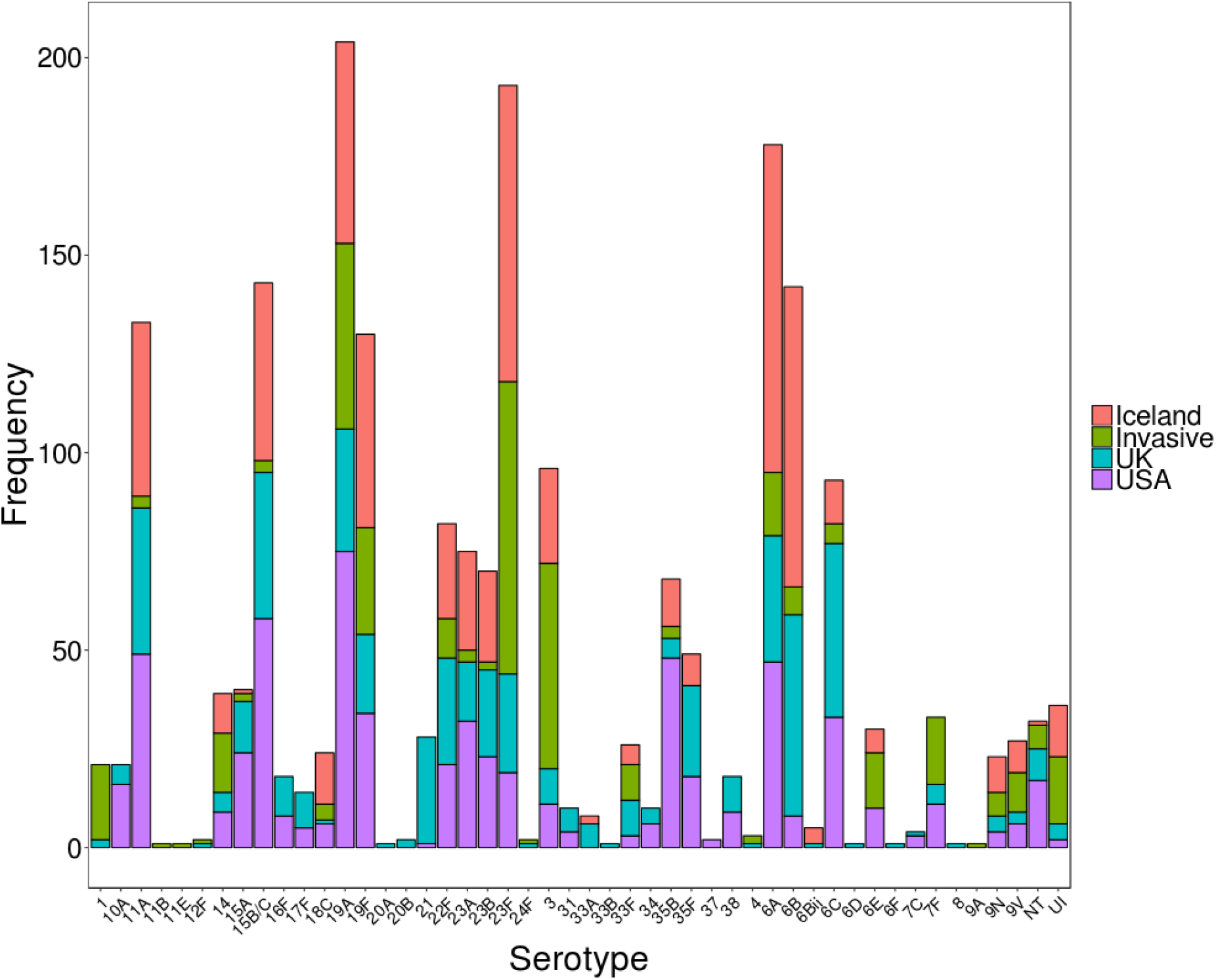
serotype distribution by dataset. Colors indicate the invasive isolates (green) and carriage isolates from Iceland (red), the UK (teal) and USA (purple). NT- non-type serotypes, UI – unidentified serotypes.

**Figure S2 (attached separately):** FasTree output for UK.

**Figure S3 (attached separately):** FasTree output for USA.

**Figure S4 (attached separately):** FasTree output for Iceland.

**Figure S5:**
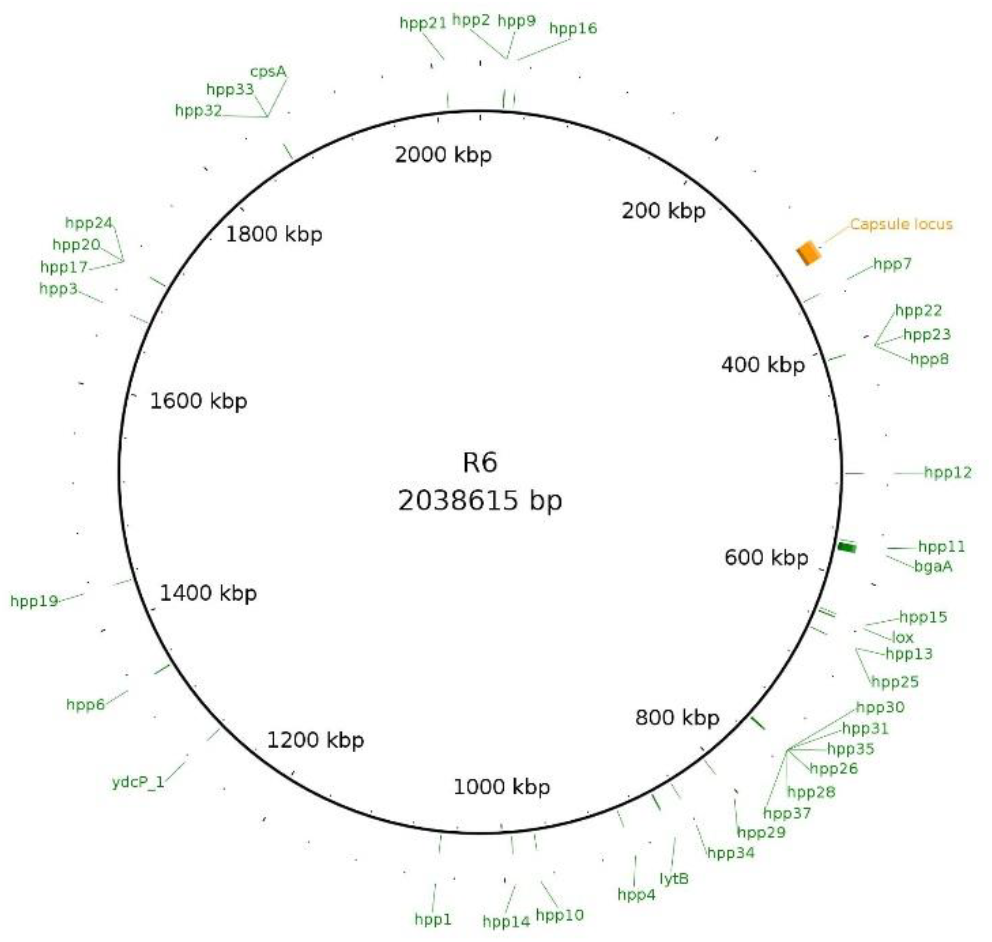
gene location plot for Isolate D39 (accession NC_008533.1).

**Figure S6:**
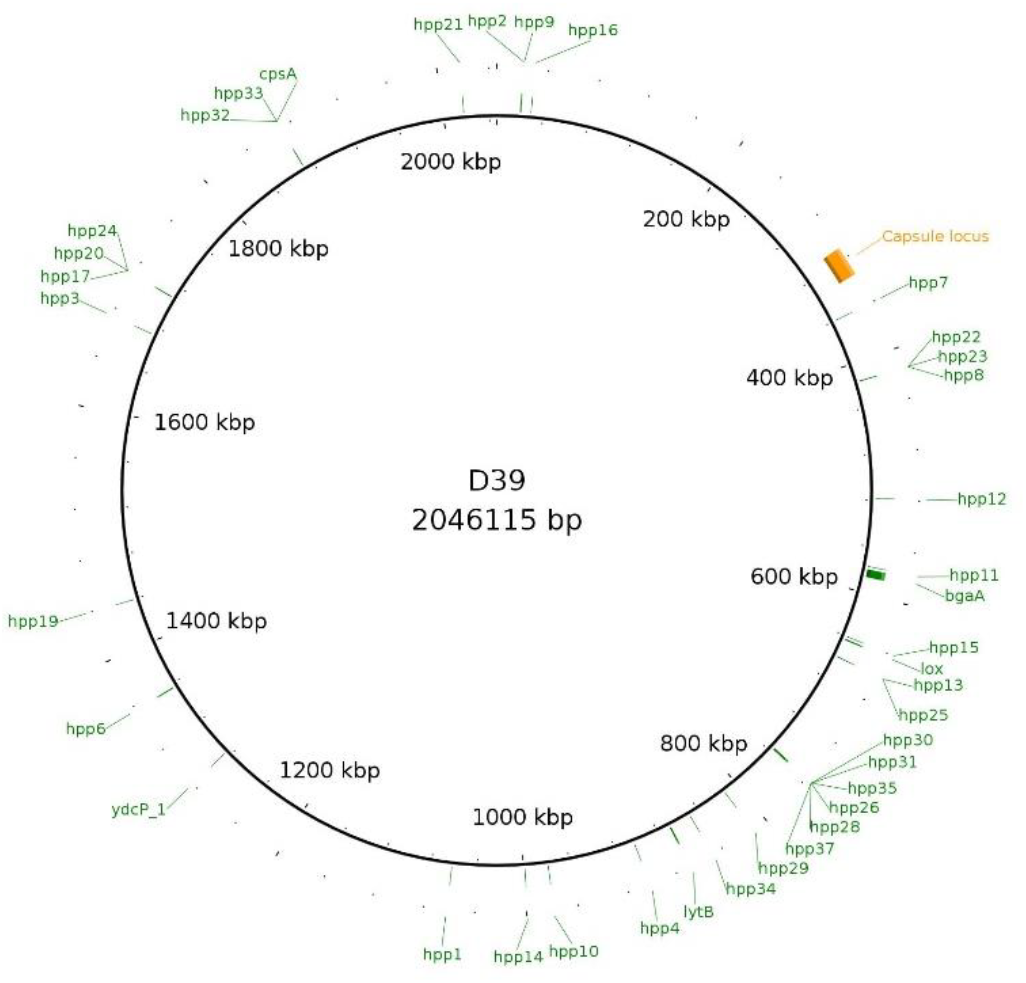
gene location plot for Isolate R6 (accession NC_003098.1).

**Figure S7:**
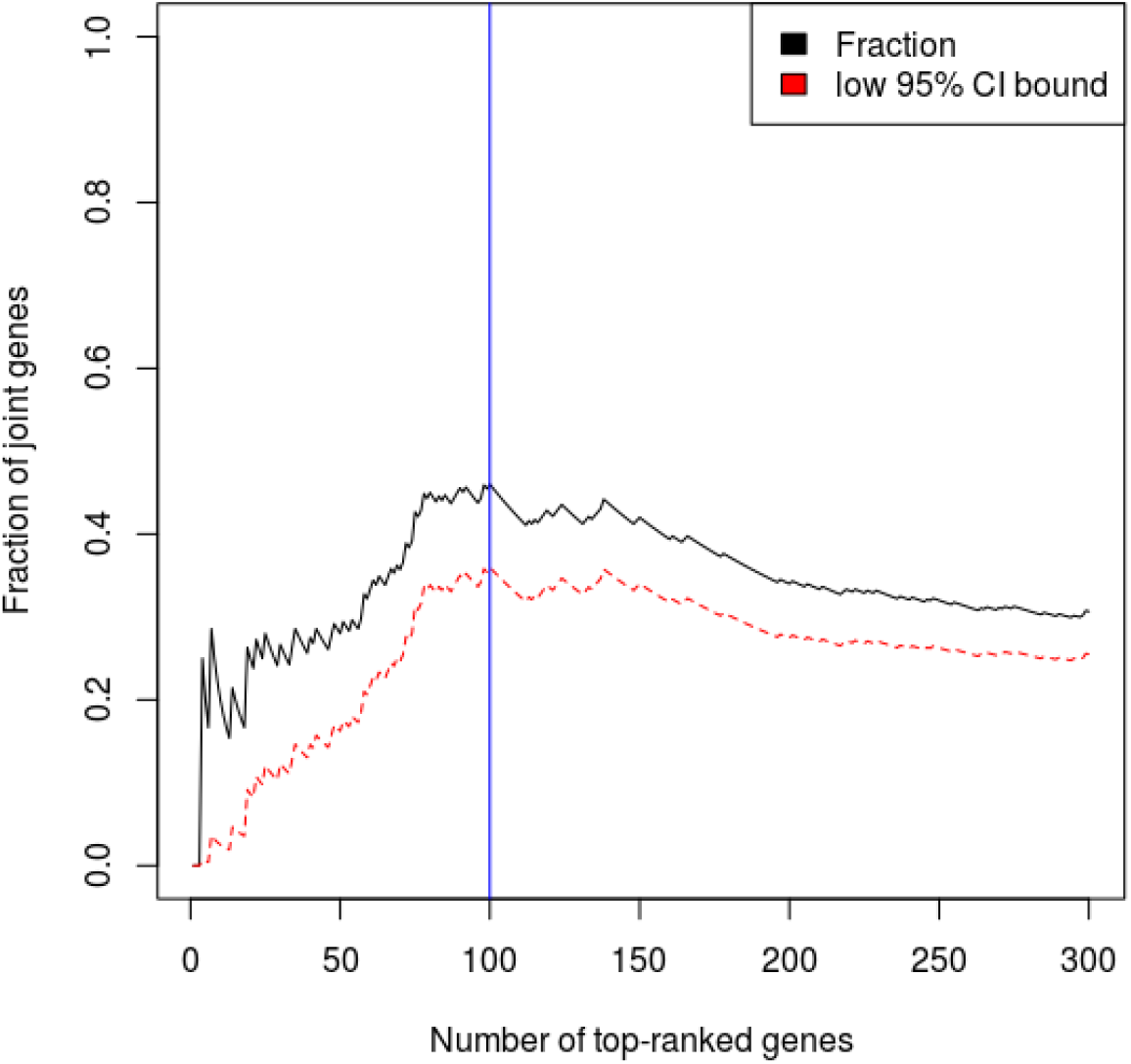
Joint gene fraction as a function of number of top-ranked genes selected. The black curve represents the fraction, while the red is the lower bound of a 95% binomial confidence interval around the fraction. The blue horizontal line indicates the number of top-ranked genes corresponding to the maximum fraction of joint genes (and the low bound of the confidence interval), at 100.

**Figure S8:**
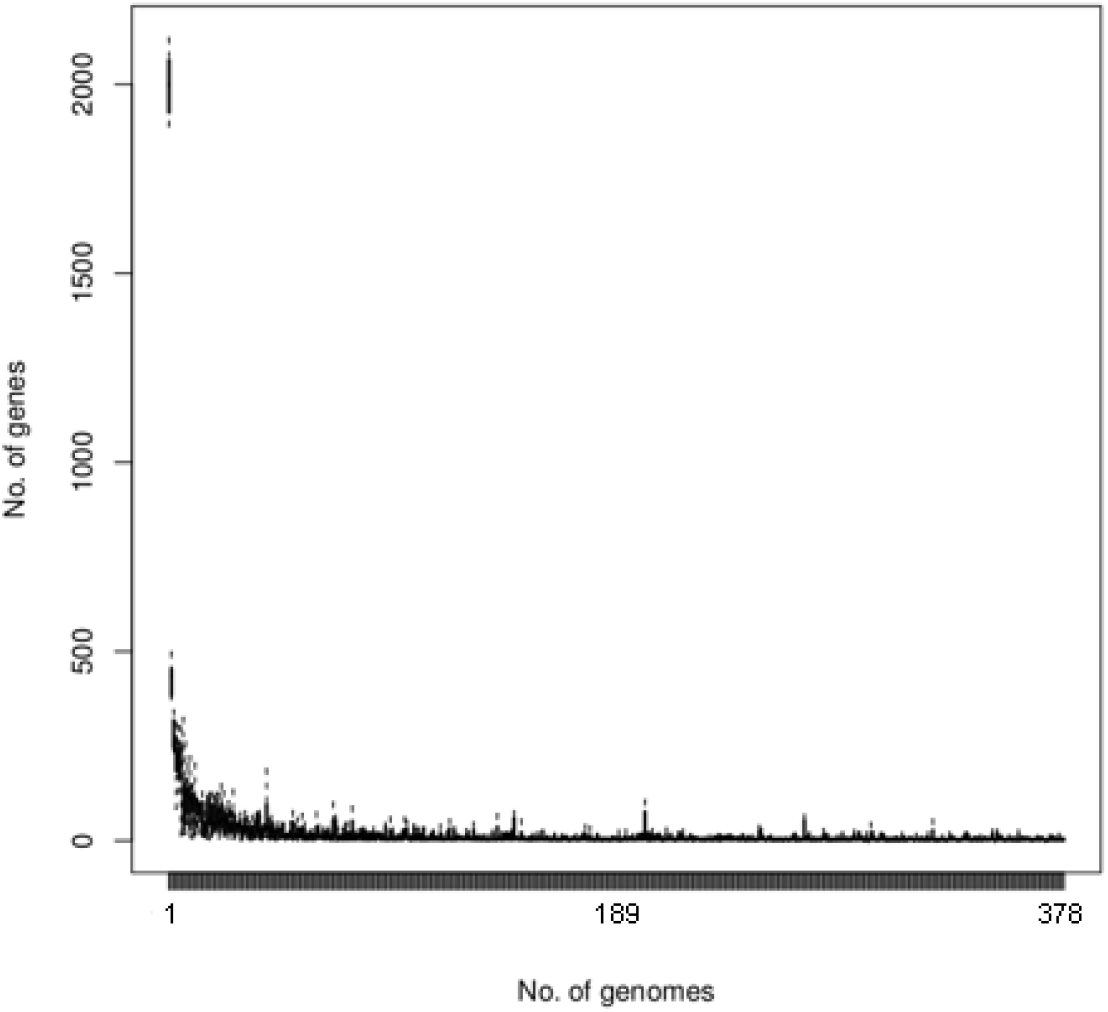
Genes added to the pangenome per new isolate. Boxplots represent the distribution of the number of new genes added to the pangenome for each additional isolate, based on 10 random samples of new isolates.

**Figure S9:**
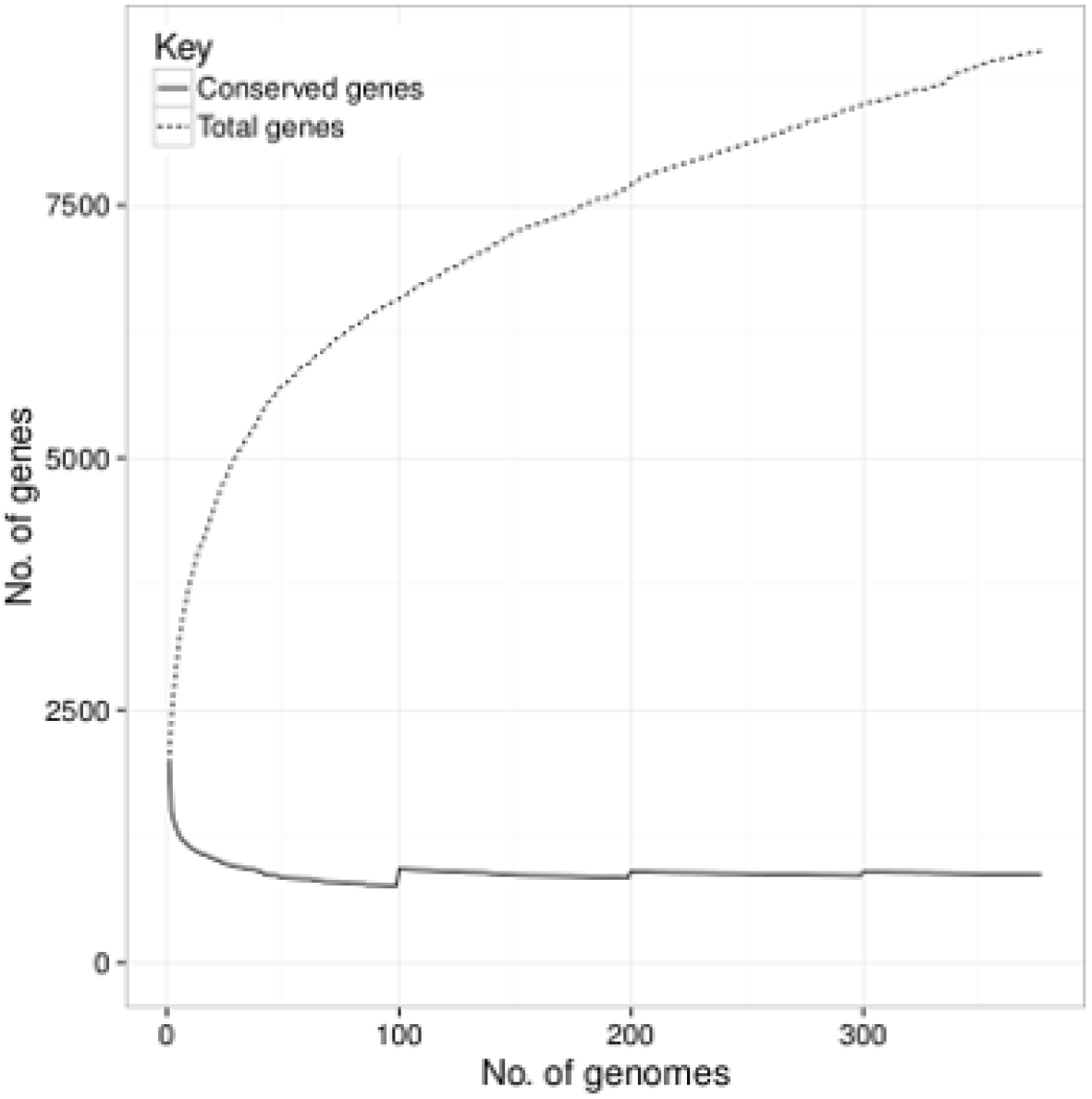
Number of core genes (present in more >99% of strains, dashed curve) and total genes per new genome (solid curve) in the invasive disease pangenome.

**Table S1 (attached separately)**: Pneumococcal sample metadata and accession numbers. Columns, from left to right, indicate the dataset from which isolate was obtained (Invasive disease, USA, UK, or Iceland), the ID number for each isolate’s genome assembly as reported on https://pubmlst.org/spneumoniae/, country of isolation, year of isolation, serotype, diagnosis of the individual carrying the isolate (bacteremia or carriage), and the ENA accession number for each isolate’s genome assembly (https://www.ebi.ac.uk/ena, where available).

**Table S2 (attached separately)**: Presence of identified genes in the invasive isolates.

**Table S3 (attached separately)**: Sequences of identified genes corresponding to Table1 in the main text.

